# Differential Mast Cell Phenotypes in Benign versus Cancer Tissues and Prostate Cancer Oncologic Outcomes

**DOI:** 10.1101/2020.07.23.216408

**Authors:** Heidi Hempel Sullivan, Janielle P. Maynard, Christopher M. Heaphy, Jiayun Lu, Angelo M. De Marzo, Tamara L. Lotan, Corinne E. Joshu, Karen S. Sfanos

## Abstract

We previously reported that high numbers of mast cells in benign (extra-tumoral) regions of the prostate are associated with worse outcomes after radical prostatectomy including biochemical recurrence and the development of metastases. Herein, on a cohort of 384 men, we performed mast cell subtyping and report that higher minimum number of the tryptase-only (MC_T_) subset of extra-tumoral mast cells is associated with increased risk of biochemical recurrence (comparing highest to lowest tertiles: HR 2.20, 95% CI 1.32-3.65; P-trend 0.004), metastases (HR 3.60, 95% CI 1.77-7.36; P-trend 0.001), and death from prostate cancer (HR 2.96, 95% CI 1.23-7.08; P-trend 0.02). RNAsequencing of benign versus cancer tissue mast cells revealed differential expression of additional site-specific genes. We demonstrate that genes more highly expressed in tumor-infiltrating mast cells, such as CXCR4 and TFE3, represent an altered tumor microenvironment. C-kit variants were also differentially expressed in benign versus cancer tissue mast cells, with C-kit variant 1 (GNNK+) mast cells identified as more prevalent in extra-tumoral regions of the prostate. Finally, using an established mouse model, we found that mast cells do not infiltrate Hi-Myc tumors, providing a model to specifically examine the role of extra-tumoral mast cells in tumorigenesis. Hi-Myc mice crossed to mast cell knockout (Wsh) mice and aged to one year revealed a higher degree of pre-invasive lesions and invasive cancer in wildtype mice versus heterozygous and knockout mice. This suggests a dosage effect where higher numbers of extra-tumoral mast cells resulted in higher cancer invasion. Overall, our studies provide further evidence for a role of extra-tumoral mast cells in driving adverse prostate cancer outcomes.

## Introduction

Mast cells are the archetypical sentinel cells of the immune system. Best characterized for their role in allergic inflammation, mast cells are distributed throughout most human tissues poised to release their pro-inflammatory granule contents upon stimulation by foreign allergens. Mast cells arise in the bone marrow, circulate as progenitors, and differentiate in their final tissue site where they remain for months or even years [1]. Since fully differentiated mast cells are not present in the circulation and only exist in limited numbers in tissues, mechanistic studies of mast cell phenotypes have proven difficult [2]. Mast cell secretory granules are armed with potent immunomodulatory components as well as proteases that serve to increase tissue and vascular permeability for immune cell infiltration. Two subtypes of mast cells have been described in humans: MC_T_ mast cells (predominant in lung) and MC_TC_ mast cells (predominant in skin). The subtypes differ in the presence of tryptases, chymases, and carboxypeptidases (MC_TC_) versus tryptases alone (MC_T_) in their secretory granules [3]. Functional differences in mast cell subtypes are less well understood, and both mast cell subtypes are present at many sites and may increase in number in relation to disease. Accordingly, the compositional and secretory functionality of mast cells in the disease microenvironment may shift [4, 5]. For example, MC_TC_ mast cells increase in number in bronchi and both MC_T_ and MC_TC_ mast cells increase in the alveolar parenchyma of patients with severe asthma [6]. The classification of mast cells into the MC_T_ and MC_TC_ phenotypes is, in fact, regarded as an oversimplification, and tissue-specific sub-populations of mast cells likely exist [3, 7]. In addition, there is mounting evidence that mast cell profiles within a given tissue microenvironment are not fixed, and may change due either to mast cell plasticity or by recruitment of mast cell precursors that eventually develop into different subtypes based on signals in the microenvironment [3, 4, 8-10].

Mast cells can mediate both innate and adaptive immune responses that, in the context of cancer, may exhibit both pro- and anti-tumorigenic effects [11, 12]. Indeed, studies examining human cancer tissues as well as using experimental models have reported both pro-tumorigenic and anti-tumorigenic functions of mast cells in a variety of cancer types (reviewed in [12]). Furthermore, mast cells may play differential, and possibly even opposing roles, in tumor initiation and progression. For example, while the majority of human cancer studies report increased numbers of mast cells in the tumor microenvironment and their potential association with tumor initiation [12], many studies report that decreased numbers or loss of mast cells within the tumor associates with a worse prognosis [13-16]. The significance of MC_T_ versus MC_TC_ mast cell subtypes in relation to cancer are even less clear. In one study, both MC_T_ and MC_TC_ cells were found to be increased in number in breast cancer, and increases in both mast cell subtypes were associated with less aggressive luminal A and luminal B molecular subtypes compared to triple-negative and HER2+ non-luminal subtypes [17]. Conversely, a lung cancer study found that low numbers of MC_TC_ mast cells were associated with a worse prognosis in a subset of stage I patients [18].

In tissue samples from specimens containing prostate cancer, there are increased mast cell numbers in areas of prostate cancer versus benign prostate tissues, and mast cells numbers are higher in density in lower Gleason grade than in higher Gleason grade cancers [14, 19-23]. We and others have reported that high intra-tumoral mast cell numbers are associated with favorable prognosis [14, 19, 23]. The opposite association exists for extra-tumoral mast cells, where high mast cell numbers are associated with worse prognosis. We previously analyzed mast cell density in relation to race and both biochemical recurrence (PSA progression) and the development of metastases after radical prostatectomy [14, 24]. Our findings highlight an important predictive role for high numbers of benign tissue (extra-tumoral) mast cell numbers in association with a higher risk of both biochemical recurrence and metastases in both African-American and European-American men [14, 24]. Previous studies have identified both MC_T_ and MC_TC_ mast cells in the prostate of men with prostate cancer [25, 26]; however, mast cell subtypes have not been previously examined in relation to race or prostate cancer prognostic factors.

In the present study, we determined if the association of mast cells with prostate cancer outcomes was attributable to a particular mast cell subtype and we further profiled RNA expression in intra-tumoral versus benign tissue mast cells. We report that high numbers of benign tissue MC_T_ mast cells specifically are associated with adverse prostate cancer oncologic outcomes after radical prostatectomy, including biochemical recurrence, metastases, and prostate cancer-specific death. We further identified differential gene expression profiles in cancer and benign tissue mast cells, including expression of C-X-C Motif Chemokine Receptor 4 (CXCR4) and Transcription Factor Binding To IGHM Enhancer 3 (TFE3) on mast cells and other immune and stromal cells within the tumor microenvironment and differential expression of KIT proto-oncogene receptor tyrosine kinase (C-kit) splice variants in cancer versus benign tissue mast cells. Finally, we examined the effects of mast cell depletion (C-kit^w-sh^ mice) and mast cell reduction (C-kit^w-sh^ heterozygous mice) on the Hi-Myc mouse prostate cancer model [27].

## Materials and Methods

### Tissue Microarray (TMA) Samples

In total, 384 men were assessed across two different cohorts. All samples were obtained and analyzed under an Institutional Review Board approved study.

The “PCBN High Grade Race” TMA set was obtained via the Prostate Cancer Biorepository Network (PCBN; http://prostatebiorepository.org). This TMA set contains radical prostatectomy cancer and benign tissue from 60 African-American men matched to 60 European-American men on age +/- 3 years, grade and stage and is enriched for cases with Gleason score ≥ 8. This 120 Case PCBN High Grade Race TMA set was designed for use in immunohistochemistry (IHC) / immunofluorescence (IF) as well as RNA *in situ* hybridization (RISH) assays, as the blocks used for TMA construction were new cases and the donor blocks as well as the TMA blocks are stored at −20 °C to preserve RNA [28]. The “Intermediate/High Risk” TMA set contains cancer and benign tissues from a radical prostatectomy cohort of primarily European-American men, retrospective case-cohort design of 356 men with intermediate or high risk disease that received no additional treatment until the time of metastases as previously described [29]; and 264 men in the sub-cohort with complete mast cell, demographic, and pathologic information available were included in the analyses. A maximum of 4 cancer spots and 4 benign tissue spots were analyzed per man.

A TMA of new cases of 5 radical prostatectomy specimens (one Gleason 6, two Gleason 7, two Gleason 8, one Gleason 9 prostate cancer) was used for the double RISH stains for C-kit/CXCR4, CXCR4/CD68, and TFE3/C-kit and for C-kit variant 1 RISH.

### Tryptase-Chymase double immunofluorescence (IF)

TMAs were treated with HTTR antigen retrieval followed by incubation with rabbit anti-chymase antibody (Abcam, Cambridge, UK, Cat No. EPR13136, 1:2000) and mouse anti-tryptase antibody (Abcam, Cambridge, UK, Cat No. Ab2378, 1:2000). Antigen detection was accomplished with goat-ant-rabbit Cy3 secondary and chicken-anti-mouse Cy5 secondary (ThermoFisher, Waltham, MA, Cat No. A11036 and A21463 respectively, both 1:100), as well as treatment with DAPI for nuclear visualization. As shown in Supplemental Figure S1A, MC_T_ cells were visualized as tryptase only positive mast cells and MC_TC_ cells were visualized as tryptase and chymase double positive mast cells.

### Scanning and analysis with TissueGnostics software

The stained IF slides were imaged with a 20X objective using the TissueFAXS Plus (TissueGnostics, Tarzana, CA) automated microscopy workstation equipped with a Zeiss Z2 Axioimager microscope. All TMAs were scanned using identical exposure times. The digitized fluorescent images were then quantified using the TissueQuest 6.0 software module to determine the number of mast cells per TMA spot (mast cell count). We compared the automated counting to manual counting of 140 TMA spots as the gold standard. We found a high correlation between manual counting and automated counting using the TissueGnostics software (R^2^ = 0.94 for total mast cells and 0.93 for MC_TC_ mast cells, Supplemental Figure S1B).

### Toluidine blue staining and laser capture microdissection (LCM)

Frozen cancer and matched benign prostate tissues from 4 radical prostatectomy specimens (all Gleason grade 9) were cut onto Leica PEN-membrane 4,0um Frame slides, fixed in 90% ethanol in DEPC, and toluidine blue stained with 0.1% toluidine blue (Sigma-Aldrich, St. Louis, MO, Cat No. 89640) in DEPC treated with 0.4U/uL Protector RNase Inhibitor (Sigma-Aldrich, St. Louis, MO, Cat No. 3335402001). LCM of toluidine blue-positive mast cells was performed on a Leica LMD 7000 Microscope (Leica, Wetzlar, Germany).

### Mast cell LCM RNA extraction and RNAseq

RNA extraction was accomplished with Qiagen RNeasy Micro Kit (Qiagen, Hilden, Germany). Cancer and benign mast cell RNA samples were pooled for each case (resulting in one pooled cancer sample and one pooled benign sample), RNA quality was assessed using a Bioanalyzer (Agilent Technologies, Santa Clara, CA), and RNA sequencing (RNAseq) was performed at the Sidney Kimmel Comprehensive Cancer Center Next Generation Sequencing Core Facility. Details of RNAseq library prep and analysis parameters are given in the Supplemental Methods.

### C-kit variant 1 quantitative reverse transcription PCR (qRT-PCR)

RNA and converted cDNA used for C-kit variant qRT-PCR was from benign and matched tumor frozen tissues (two Gleason 6, five Gleason 7, one Gleason 8, and two Gleason 9 prostate cancers) from radical prostatectomy specimens as previously described [30]. qRT-PCR was performed with C-kit variant 1 forward primer 5’-CAACAAAGAGCAAATCCATCCC-3’ and reverse primer 5’-CATCACAATAATGCACATCATGCC-3’ with iQ SYBR Green supermix (BioRad, Hercules, CA, Cat No. 1708882). The PCR program was as follows: 94°C for 3 min followed by 40 cycles of 94°C for 30 seconds, 50°C for 30 seconds and 72°C for 30 seconds. Each sample was normalized to GAPDH as previously described [30].

### Control cell plugs for C-kit V1 and V2, CXCR4, TFE3

Control cell plugs were made by transfecting PC3 cells with C-kit Variant 1 human ORF cDNA clone (Origene, Rockville, MD, Cat No. SC323577, NM_000222), C-kit Variant 2 human ORF cDNA clone (Origene, Rockville, MD, Cat No. SC316285, NM_001093772), CXCR4 human ORF cDNA clone (Origene, Rockville, MD, Cat No. SC117951, NM_003467), TFE3 variant 2 human ORF cDNA clone (Origene, Rockville, MD, Cat No. SC336310, NM_001282142). Transfection was accomplished with Lipofectamine 2000 (Life Technologies, Carlsbad, CA, Cat No. L0000-001). Cells were spun down, fixed in 10% formalin for 48hrs, and paraffin embedded.

### RNA in situ hybridization (RISH)

All positive and negative controls for RISH are shown in Supplemental Figure S2. 1zz probes were designed by Advanced Cell Diagnostics (ACD, Newark, CA) to be specific to C-kit variant 1 but not variant 2 using the Basescope manual red kit (Supplemental Figure S2A). Staining of control plugs was also done with DNAse I (Sigma-Aldrich, St. Louis, MO, Cat No. D5319) or RNAse A (DNAse-free Affymetrix, Santa Clara, CA, Cat No. 78020Y) treatment steps to verify detection of RNA versus genomic DNA (data not shown). CXCR4 probe (ACD, Newark, CA, Cat No. 310511) and TFE3 probe (ACD, Newark, CA, Cat No. 430461) were used with the ACD RNAscope 2.5 manual brown kit. ACD Dual RNAscope manual kit was used to double stain for C-kit (Cat No. 606401)/CXCR4 (Cat No. 310511-C2), CXCR4 (Cat No. 310511)/CD68 (Cat No. 560591-C2), and TFE3 (Cat No. 430461)/C-kit (Cat No. 606401-C2) (ACD, Newark, CA).

### Hi-Myc/Wsh mouse generation

Mast cell deficient C-kit^w-sh^ (Wsh) mice were obtained from Jackson Laboratory (Stock No. 012861). Hi-Myc mice were obtained from Frederick National Laboratory (Strain No. 01XK8) [27]. Hi-Myc mice were crossed with Wsh mice to the second generation. Littermates of the following genotypes were aged for 1 year (360 ± 13 days): Hi-Myc +/-, Kit+/+ (WT), Hi-Myc +/-, Kit+/- (HT), Hi-Myc +/-, Kit-/- (KO), n=10 of each genotype. All mice were second generation and littermates in order to control for mixed background.

### Slide staining, annotation, and analysis

FFPE slides of mouse prostate (all lobes) were hematoxylin and eosin stained. Adjacent slides were also stained with toluidine blue (1% toluidine blue in 1% sodium chloride and 7% ethanol) for visualization and quantification of mast cells by lobe. Slides were scanned at 20X using the Aperio ScanScope (CS model, Aperio, Vista, CA) and annotated (blinded to group status) in Aperio Imagescope (version 12.2.2.8013) for prostatic intraepithelial neoplasia (PIN), cribriform PIN, and invasive cancer area by prostate lobe [31]. Toluidine blue-positive mast cells were counted manually using the Aperio Imagescope software. For CXCR4 image analysis, slides were scanned using the Aperio ScanScope then viewed and analyzed using Aperio ImageScope Software. TMA scanned images were segmented then analyzed using TMAJ and FrIDA software (version 3.15.0) as previously described [28].

### Statistical analysis

In our previous study [14], the minimum number of mast cells counted among a man’s TMA spots in tumor and benign tissue was the most robustly different between PSA recurrence cases and controls. Therefore, similar to our previous analysis, all men were categorized into tertiles of minimum mast cell count. The minimum mast cell count is the lowest number of mast cells counted among up to 4 benign or 4 cancer tissue spots. Cox proportional hazards regression was used to estimate the hazard ratio and 95% CI of biochemical recurrence and metastases by tertile of minimum mast cell count. All analyses were adjusted for age, race (European-American vs. African-American), grade (prostatectomy Gleason sum <=6, 3+4, 4+3, >=8), stage (<=T2 vs. >T2), PSA (continuous) and BMI (continuous). P-for trend was estimated by entering a continuous variable for each tertile in the model. All tests were 2-sided, with p<0.05 considered to be statistically significant. All were conducted using SAS 9.4 (SAS Institute).

For all other analyses, data were compared between groups by two-tailed Mann Whitney U or Wilcoxon matched-pairs signed rank test using GraphPad Prism Software (version 7.01; GraphPad Software, Inc., San Diego, CA). Values were considered significantly significant at p<0.05.

## Results

### Mast cell subtypes in prostatectomy specimens

We used a dual IF stain for tryptase and chymase to analyze the 120 Case PCBN High Grade Race TMA set to assess mast cell subtypes between cancer and benign tissues in African-American and European-American men. We used automated image analysis to identify and count the number of MC_T_ and MC_TC_ in each tissue spot represented across the TMAs (Supplemental Figure S1). Both mast cell subtypes were present in both cancer and benign prostate tissues (Figure 1A, B). MC_T_ mast cells were more abundant than MC_TC_ mast cells in both benign (median 8 versus 4.5 cells per tissue spot, p<0.0001, Mann-Whitney test) and cancer (median 9 versus 3 cells per tissue spot, p<0.0001) tissues (Figure 1B). The same trend of significantly lower MC_TC_ mast cells counts in both benign and cancer tissues was observed for both African-American and European-American men (Figure 1C). There were higher numbers of MC_TC_ mast cells in benign versus cancer tissue spots in African-American men (p = 0.048) and higher numbers of MC_T_ mast cells in benign tissues of European-American men versus benign tissues of African-American men (p = 0.034, Figure 1C).

**Figure 1.**
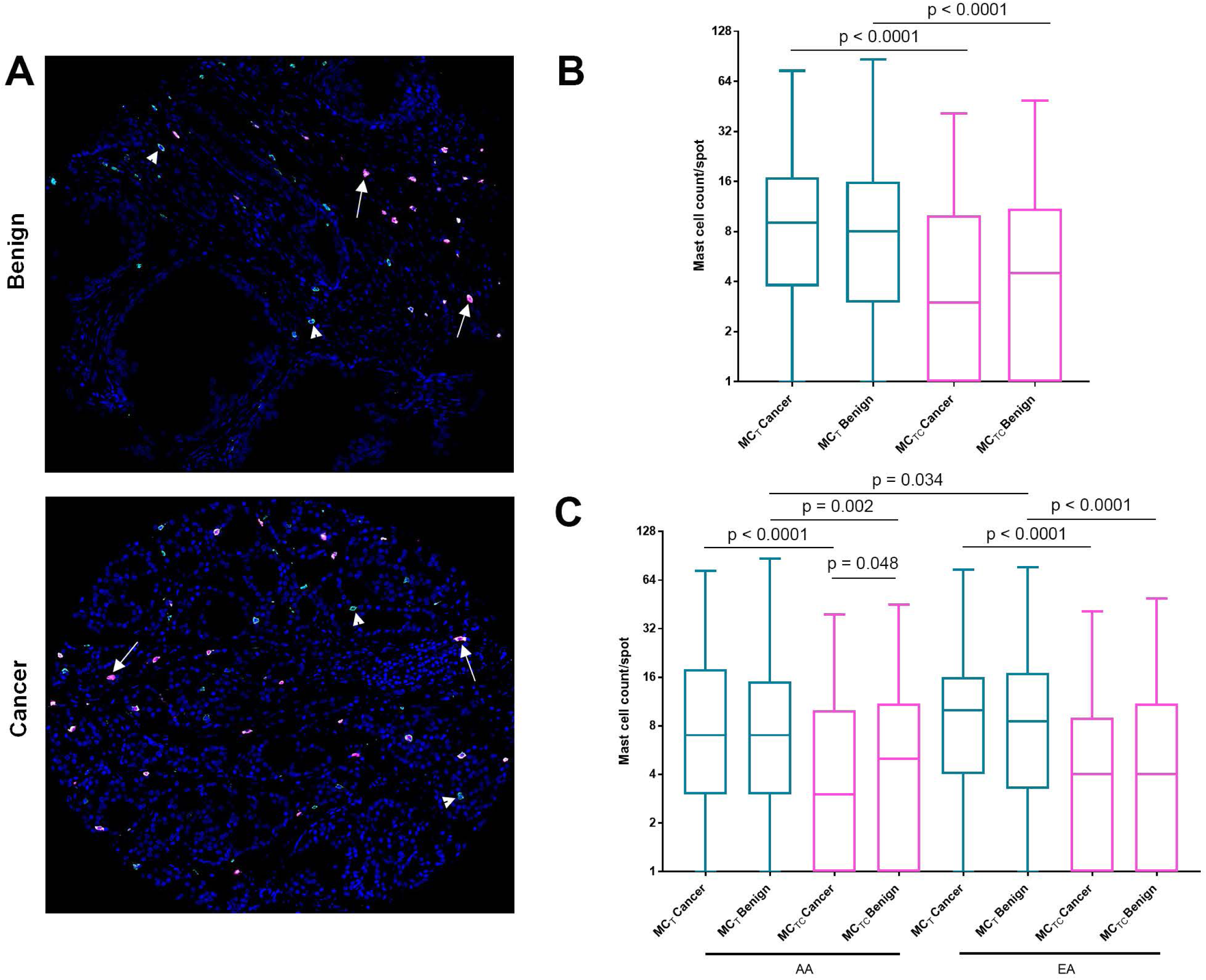
Both MC_T_ and MC_TC_ mast cell subtypes are present in benign and cancer regions of radical prostatectomy specimens. A) Example of tryptase positive MC_T_ mast cells (green, arrowheads) and tryptase+chymase positive MC_TC_ mast cells (red, arrows) in benign and cancer TMA tissue spots. B) Log2 of mast cell count per tissue spot of each mast cell subtype in benign and cancer spots. C) Log2 of mast cell count per tissue spot of each mast cell subtype in benign and cancer spots in African-American versus European-American men.

### Mast cell subtypes and oncologic outcomes

We next assessed MC_T_ and MC_TC_ mast cells in the Intermediate/High Risk TMA set, which was designed to examine biomarkers in relation to prostate cancer outcomes including biochemical recurrence, metastasis, and prostate cancer-related death [29]. Similar to the PCBN High Grade Race TMA, we observed lower MC_TC_ mast cell counts than MC_T_ mast cells in both benign (median 7 versus 3 cells per tissue spot, p<0.0001, Mann-Whitney test) and cancer tissues (median 7 versus 4 cells per tissue spot, p<0.0001). In this TMA set, there were significantly higher MC_TC_ mast cell counts in cancer versus benign tissues (median 4 versus 3 cells per tissue spot, p<0.0001, Supplemental Figure S3).

In our previous studies, the minimum number of mast cells counted among a man’s TMA spots in cancer and benign tissue was the most robustly different between PSA recurrence cases and controls [14] and metastasis cases and controls [24]. Therefore, similar to our previous analysis, all men were categorized into tertiles of minimum mast cell count. As expected based on our previous study [24], as compared the first tertile of minimum mast cell count (inclusive of all mast cells) for benign tissue, the risk of prostate cancer biochemical recurrence and metastasis significantly increased with increasing tertiles of minimum mast cell count (p-trend=0.03 and 0.01, respectively, Table 1). Of interest, we additionally observed a significantly increased risk of prostate cancer-related death with higher minimum numbers of benign tissue mast cells (p-trend=0.02).

**Table 1.**
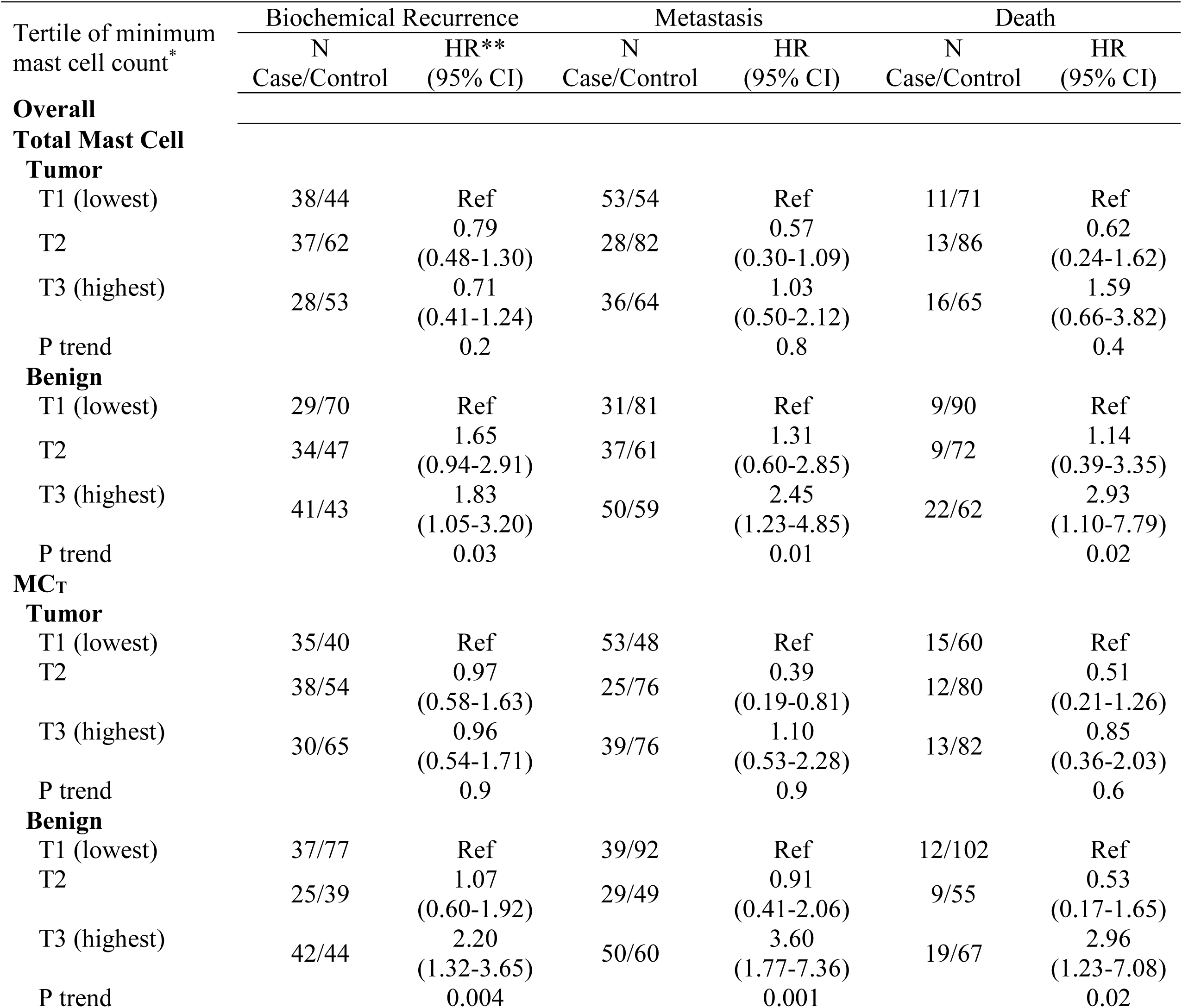
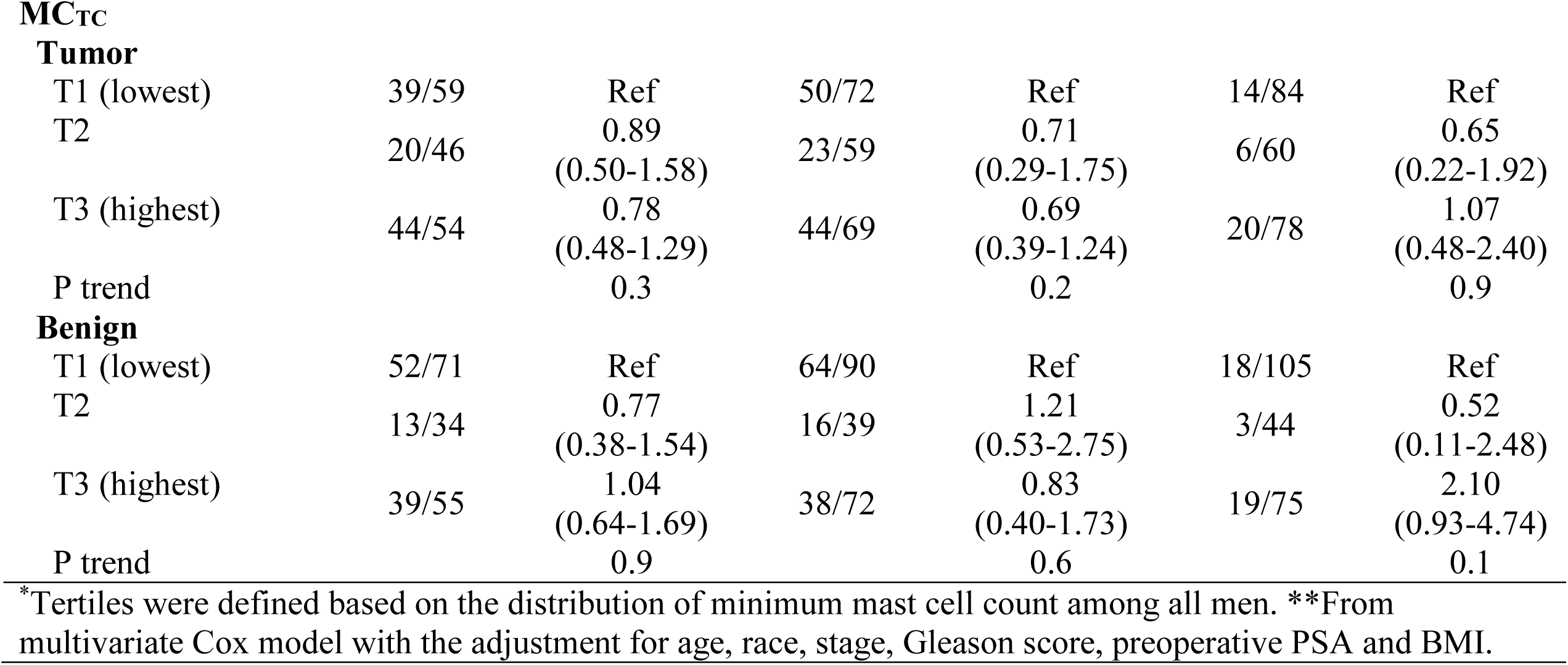
Association between tertile of minimum mast cell count and biochemical recurrence, prostate cancer metastases, and prostate cancer death.

When evaluated by mast cell subtype, the association between high benign tissue mast cells numbers and adverse prostate cancer outcomes was only present among MC_T_ mast cells. Men in the third tertile of minimum mast cell count for benign tissue had a 2.2-fold increased risk of biochemical recurrence (p-trend=0.004), 3.6-fold increased risk of metastases (p-trend=0.001), and 2.96-fold risk of prostate cancer-related death as compared to men in the first tertile (p-trend=0.02, Table 1). In contrast, there was no association between high benign tissue MC_TC_ numbers and adverse prostate cancer outcomes.

### Phenotyping of benign versus tumor-infiltrating mast cells by RNAseq

In order to further examine phenotypic differences between benign tissue versus prostate tumor-infiltrating mast cells, we devised a strategy whereby we could isolate mast cells from these distinct regions using LCM and toluidine blue stain (a mast-cell-specific stain that does not interfere with LCM or RNA extraction, Supplemental Figure S4). Laser captured mast cells from cancer and benign regions (n = 200-400 cells per case) from four high grade (Gleason grade 9, Grade group 5) prostate cancer cases were pooled and extracted RNA was submitted for RNAseq. The results of these analyses indicated that mast cells were successfully recovered from the tissues, as many known mast cell markers, including tryptase (TPSAB1, TPSB2), chymase (CMA1), mast cell carboxypeptidase A (CPA3), and IgE receptor (FCER1A, FCER1G), were represented in the RNAseq data (Figure 2, Supplemental Table S1).

**Figure 2.**
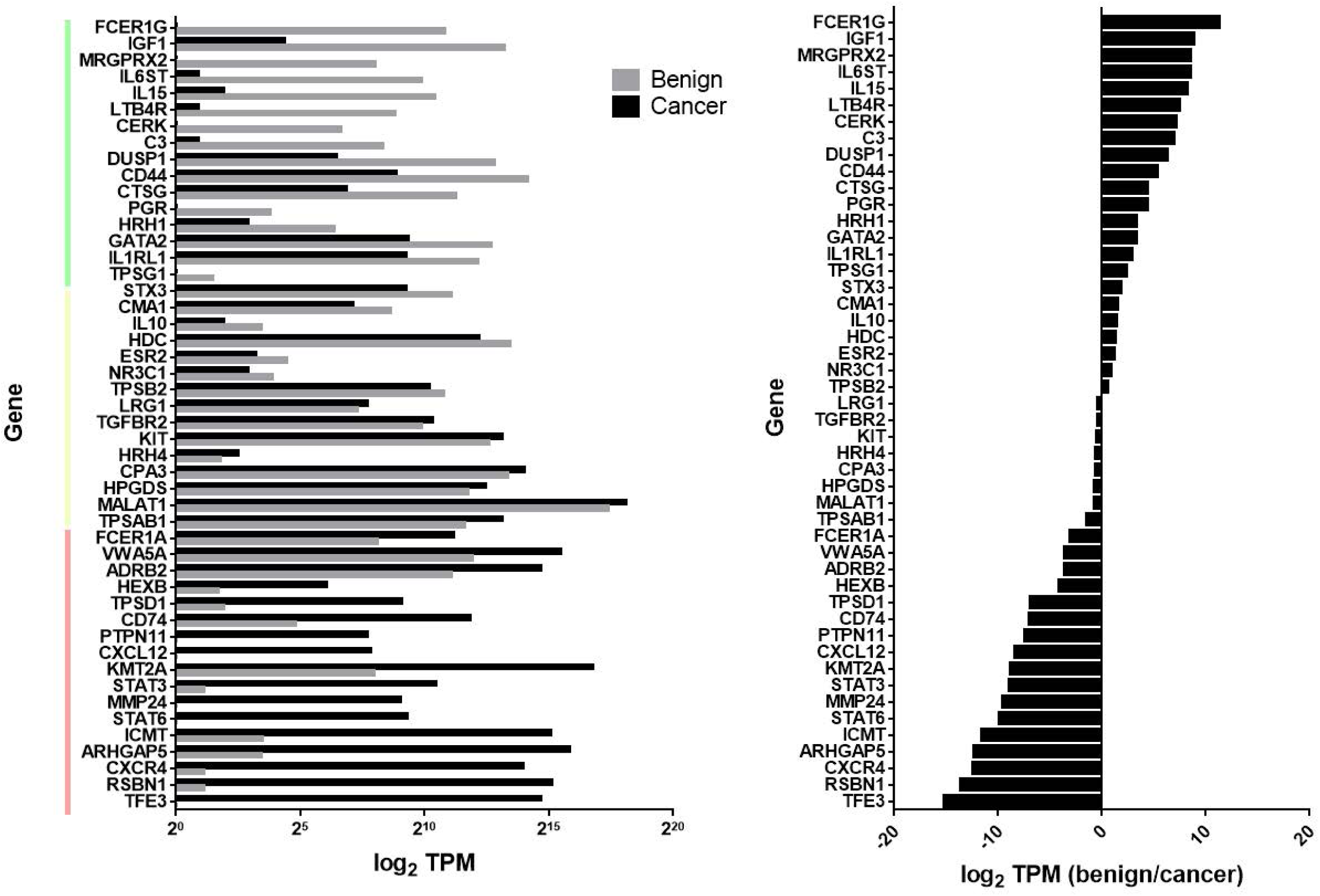
Differential gene expression in benign versus cancer tissue mast cells. Left, transcripts per million (TPM, log_2_) of genes more highly expressed in benign tissue mast cells (green bar), roughly equivalent in benign and cancer tissue mast cells (yellow bar), or more highly expressed in cancer tissue mast cells (red bar). Right, TPM (log_2_) of genes expressed in benign tissue mast cells relative to cancer tissue mast cells.

### Mast cell markers represent an altered prostate tumor microenvironment

We noted with interest the higher expression of CXCR4 on mast cells isolated from cancer regions (Figure 2). CXCR4 is expressed on some mature mast cells as well as mast cell progenitors that may be recruited to tissues via expression of its receptor CXCL12 (also known as stromal cell-derived factor 1 or SDF-1) [32, 33]. We further evaluated expression of CXCR4 on prostate tumor-infiltrating mast cells using a dual-RISH stain for CXCR4 and C-kit on a 5 case TMA. These analyses revealed that some prostate tumor-infiltrating mast cells do express CXCR4, although there were many additional CXCR4-positive cells in the tumor microenvironment that were not mast cells (Figure 3A-B). We further performed a dual-RISH stain for CXCR4 and CD68 to identify additional tumor-infiltrating CXCR4-positive cells as macrophages (Figure 3C). Likewise, lymphocyte clusters near cancer were CXCR4-positive (Figure 3D). Mast cells in benign/stromal regions of the prostate were generally CXCR4-negative (Figure 3E-F).

**Figure 3.**
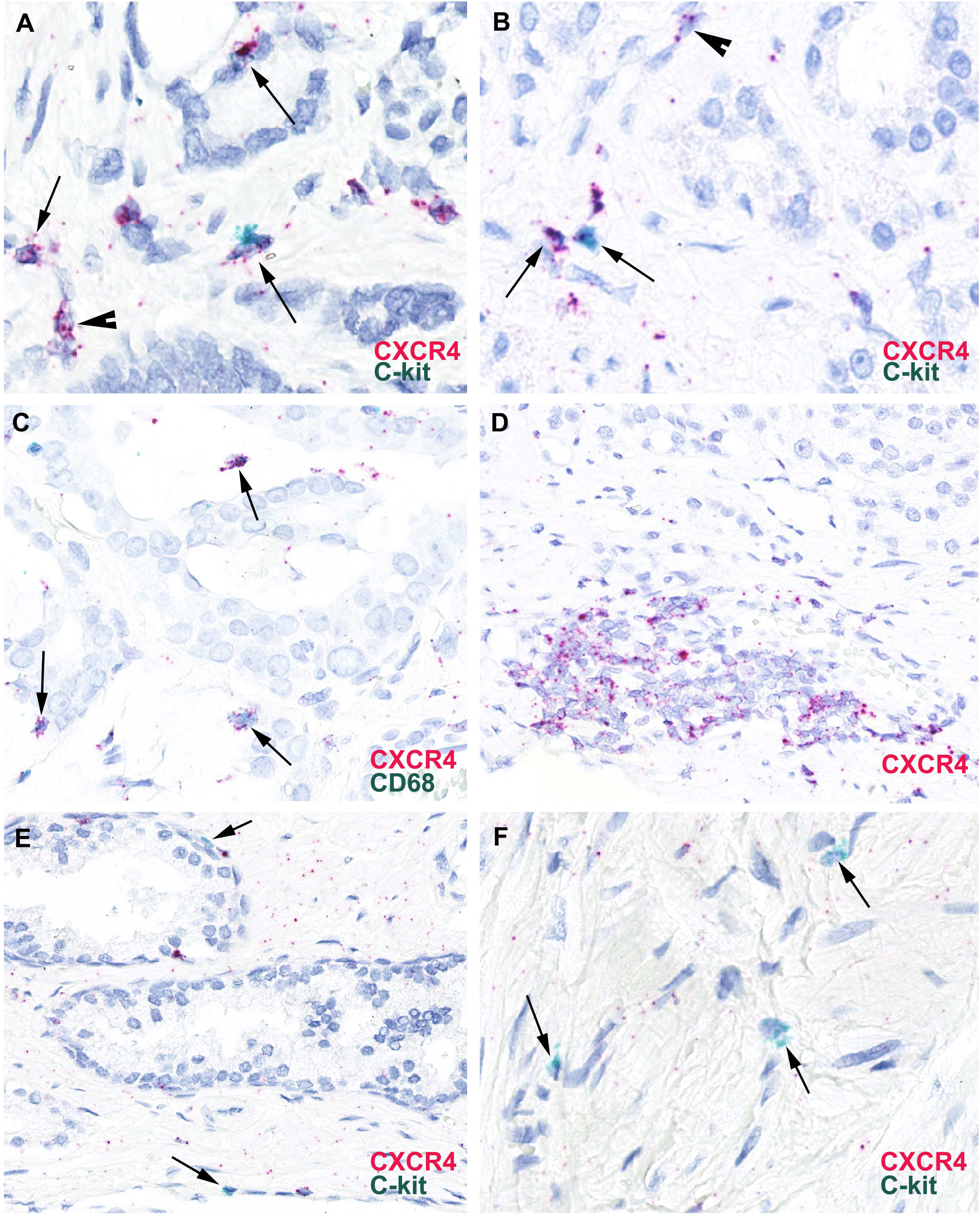
CXCR4 expression on immune cells in the prostate tumor microenvironment. A-B) Dual-RISH stain for CXCR4 (red) and C-kit (mast cell marker, green). Arrows denote dual CXCR4/C-kit positive mast cells in cancer regions. Arrowheads denote CXCR4 positive cells that are not mast cells. C) Dual-RISH stain for CXCR4 (red) and CD68 (macrophage marker, green). Arrows denote dual CXCR4/CD68 positive macrophages in a cancer region. D) CXCR4 positive (red) lymphocytes in a cancer region. E-E) Dual-RISH stain for CXCR4 (red) and C-kit (green) in benign tissue regions. Arrows denote C-kit positive (green) but CXCR4 negative mast cells.

Due to the apparent increased expression of CXCR4 on immune cells within the prostate tumor microenvironment compared to benign tissues, we further assessed CXCR4 expression with RISH on the 120 Case PCBN High Grade Race TMA set, which was designed with considerations for RNA preservation for use in CISH assays [28]. These analyses confirmed differential expression of CXCR4 in cells within the stromal compartment in cancer versus benign tissues (Figure 4A). When quantified by image analysis, CXCR4 expression was significantly higher in cancer versus benign tissue spots (p = 0.012, Wilcoxon signed-rank test, Figure 4B). Furthermore, CXCR4 expression was higher in both benign and cancer tissues in higher grade (Grade group 3-5) versus lower grade (Grade group 1-2) cases (p = 0.012 and p < 0.0001, Wilcoxon signed-rank test, respectively, Figure 4C).

**Figure 4.**
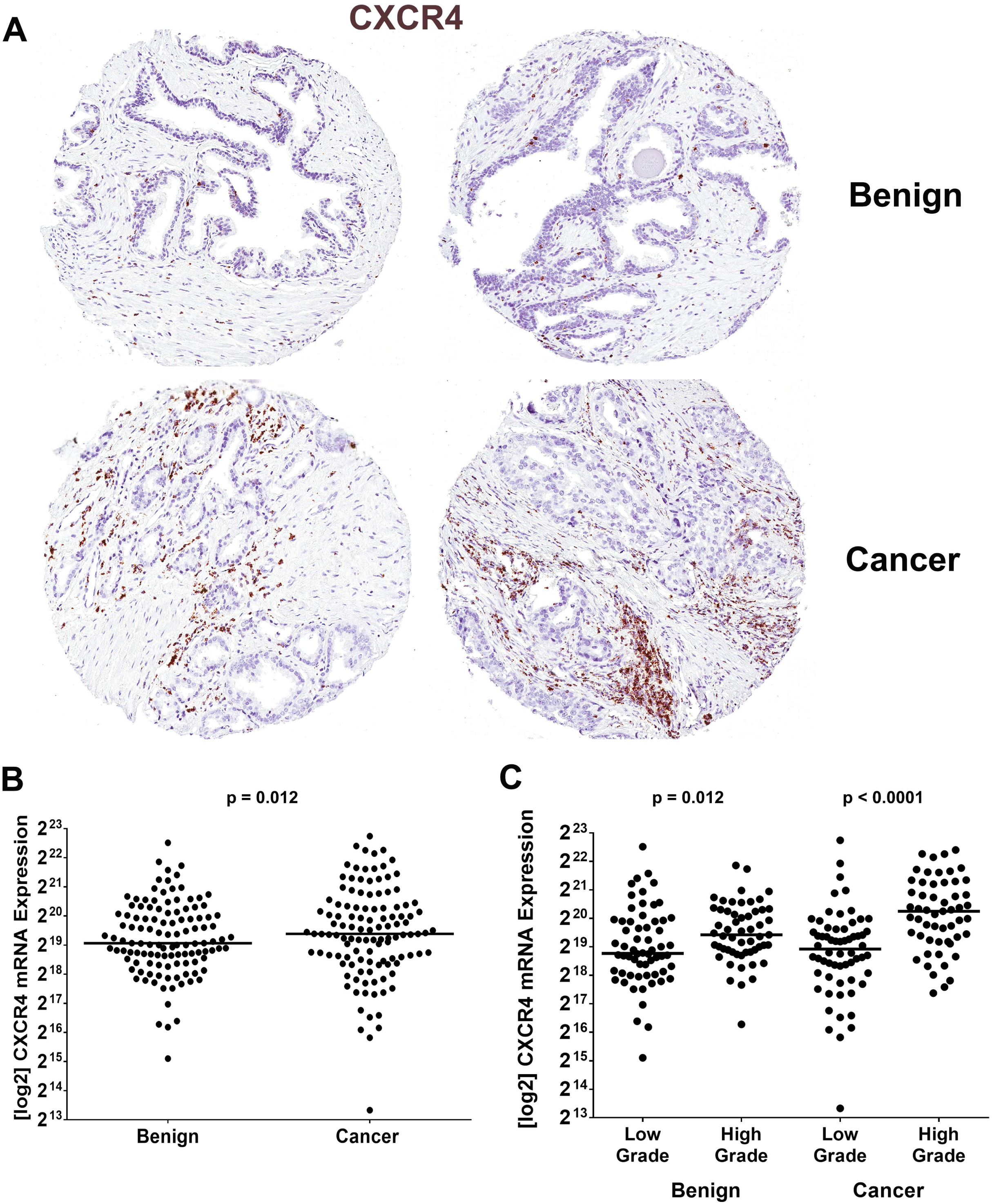
CXCR4 quantification in the 120 Case PCBN High Grade Race TMA set. A) TMA spots of benign versus cancer tissues stained with RISH for CXCR4. Note markedly increased presence of CXCR4 positive cells in the stromal compartment of the cancer spots. B) Log_2_ of median CXCR4 expression per TMA spot per case in benign versus cancer tissues. C) Log_2_ of median CXCR4 expression per TMA spot per case in benign versus cancer tissues in low (Grade group 1-2) versus higher grade (Grade group 3-5) cases.

When the benign tissue spots were further characterized as to whether they contained atrophy or PIN, we noted that CXCR4 expression was significantly increased in regions containing prostatic atrophy compared to benign, non-atrophic tissues (p < 0.0001) and PIN (p = 0.046), but not cancer tissues (Mann-Whitney test, Supplemental Figure S5A). Increased CXCR4 in prostatic atrophy was associated with increased immune cell infiltration of these areas. There was an increased median CXCR4 expression in benign tissues from higher grade versus lower grade cases from both African-American and European-American men, but this was only significant for African-American men (p = 0.034, Mann-Whitney test, Supplemental Figure S5B). Similarly, there was an increased median CXCR4 expression in cancer tissues from higher grade versus lower grade cases from both African-American and European-American men, but this was only significant for European-American men (p < 0.0001, Mann-Whitney test, Supplemental Figure S5C).

We similarly noted with interest markedly increased expression of TFE3 on the mast cell sample from cancer regions (Figure 2). TFE3 is a transcription factor and regulator of the alpha subunit of the high-affinity IgE receptor (FcεRIα, also found to be increased in mast cells isolated from cancer regions, Figure 2), associates with the C-kit promoter upon mast cell activation, and is upregulated following mast cell immunologic triggering [34]. We further evaluated expression of TFE3 on prostate tumor-infiltrating mast cells using a dual-RISH stain for TFE3 and C-kit on the 5 case TMA. These analyses demonstrated that TFE3 is expressed in both benign and cancer epithelial cells in the prostate (Figure 5A, B). Although some tumor-infiltrating mast cells were found to be double positive for C-kit and TFE3 (Figure 5C,D) and generally mast cells in benign regions were not (Figure 5B), we could not rule out that the increased TFE3 expression in the sample of mast cells isolated from cancer was not a factor of their proximity to the TFE3 expressing cancer cells. Whereas we carefully targeted mast cells in our LCM assay, it is possible that we obtained some cells immediately adjacent to the mast cells (Supplemental Figure S4). Indeed, a known prostate cancer marker, Alpha-Methylacyl-CoA Racemase (AMACR), was also highly expressed in the mast cell sample isolated from cancer tissues, likely indicating the presence of RNA captured from cancer cells (Supplemental Table S2). Unlike the tumor-infiltrating mast cells (Figure 5C,D), mast cells present in benign tissues are typically in stromal regions not immediately adjacent to the epithelial cells (Figure 5B).

**Figure 5.**
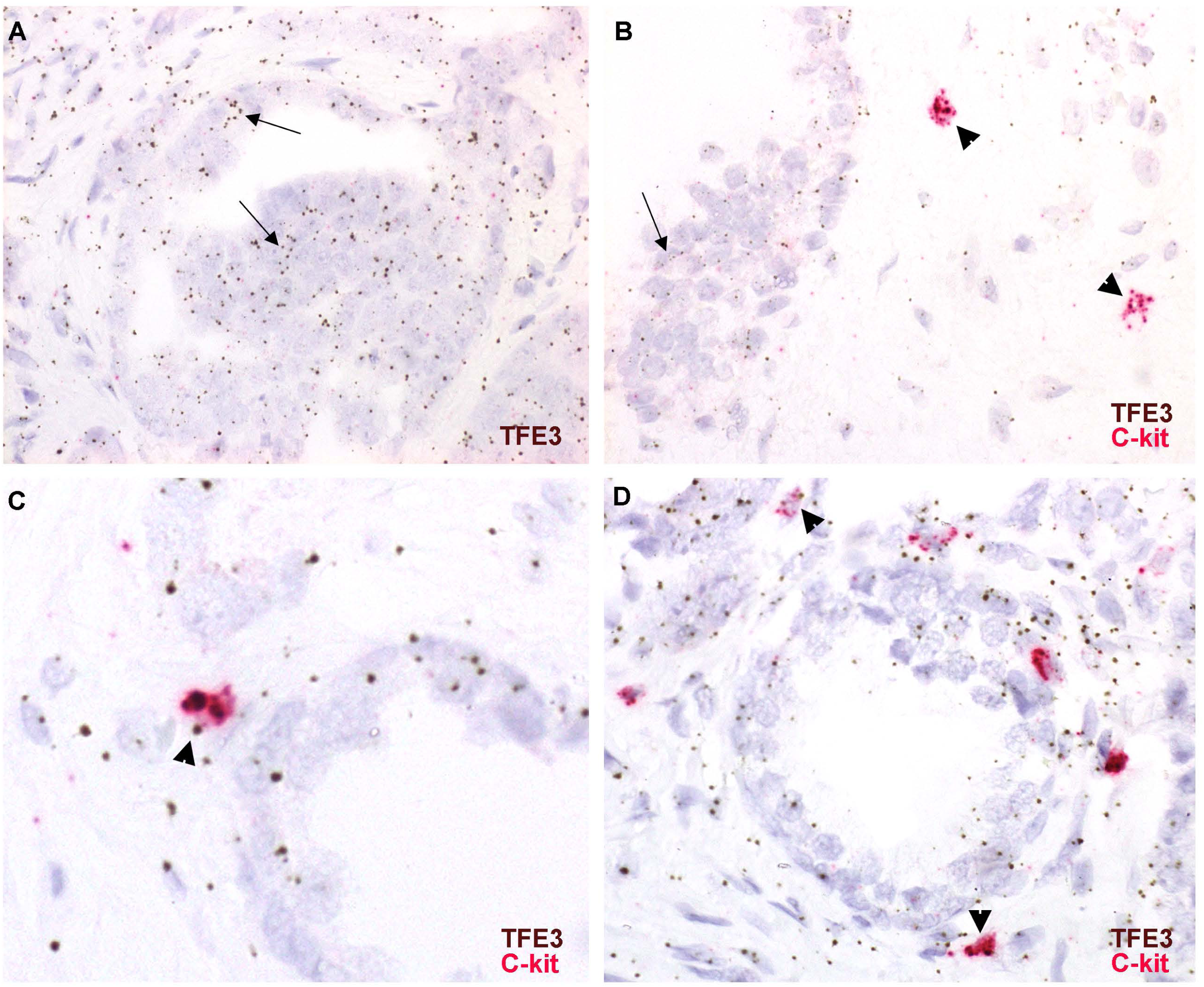
TFE3 expression on prostate epithelial cells and mast cells. A) TFE3 expression in epithelial cells in an area of prostate cancer (arrows). B) TFE3 expression in benign prostate epithelial cells (arrow). C-kit positive mast cells are negative for TFE3 in this benign region (arrowheads). C,D) TFE3 expression on C-kit positive mast cells in cancer (arrowheads).

### Reciprocal expression of C-kit receptor splice variants in benign versus tumor tissue mast cells

Although overall C-kit expression was roughly equivalent between mast cells isolated from cancer and benign tissues (Figure 2), we noted differential expression of C-kit splice variants, with Variant 1 (V1) showing higher expression in benign tissue mast cells, and Variant 2 (V2) showing higher expression in cancer tissue mast cells (Supplemental Table S3). C-kit V2 differs from V1 by the absence via an in-frame splice site variation of 12 base pairs corresponding to the amino acids GNNK in the juxtamembrane domain, and functional differences in the isoforms have been described [35, 36]. Thus, we confirmed the differential expression of the C-kit V1 transcript in benign versus cancer tissue mast cells due to our finding that benign tissue mast cells specifically are associated with adverse prostate cancer outcomes (Table 1). We designed an ACD Basescope RISH assay specific for C-kit V1 that required the presence of the 12 bp region that is absent from V2 for hybridization of the probe pairs (Supplemental Figure S2A).

Only a subset of mast cells were found to express the C-kit V1 variant in the 5 case TMA, however these V1-expressing cells were observed in association with (and interesting generally in close proximity to) benign versus cancer glands (Figure 6A). We further assessed differential expression of the C-kit variants with qRT-PCR using primers designed against the C-kit variant 1 region that is absent in variant 2 on RNA collected from prostate tumor and matched benign tissues. The RNA samples were from harvested frozen prostate tissues (not LCM isolated mast cells) from a range of Gleason score tumors and their adjacent benign prostate tissues. In 8 of 10 cases, expression of C-kit V1 was higher in RNA from benign tissues than in the matched cancer tissue RNA (Figure 6B).

**Figure 6.**
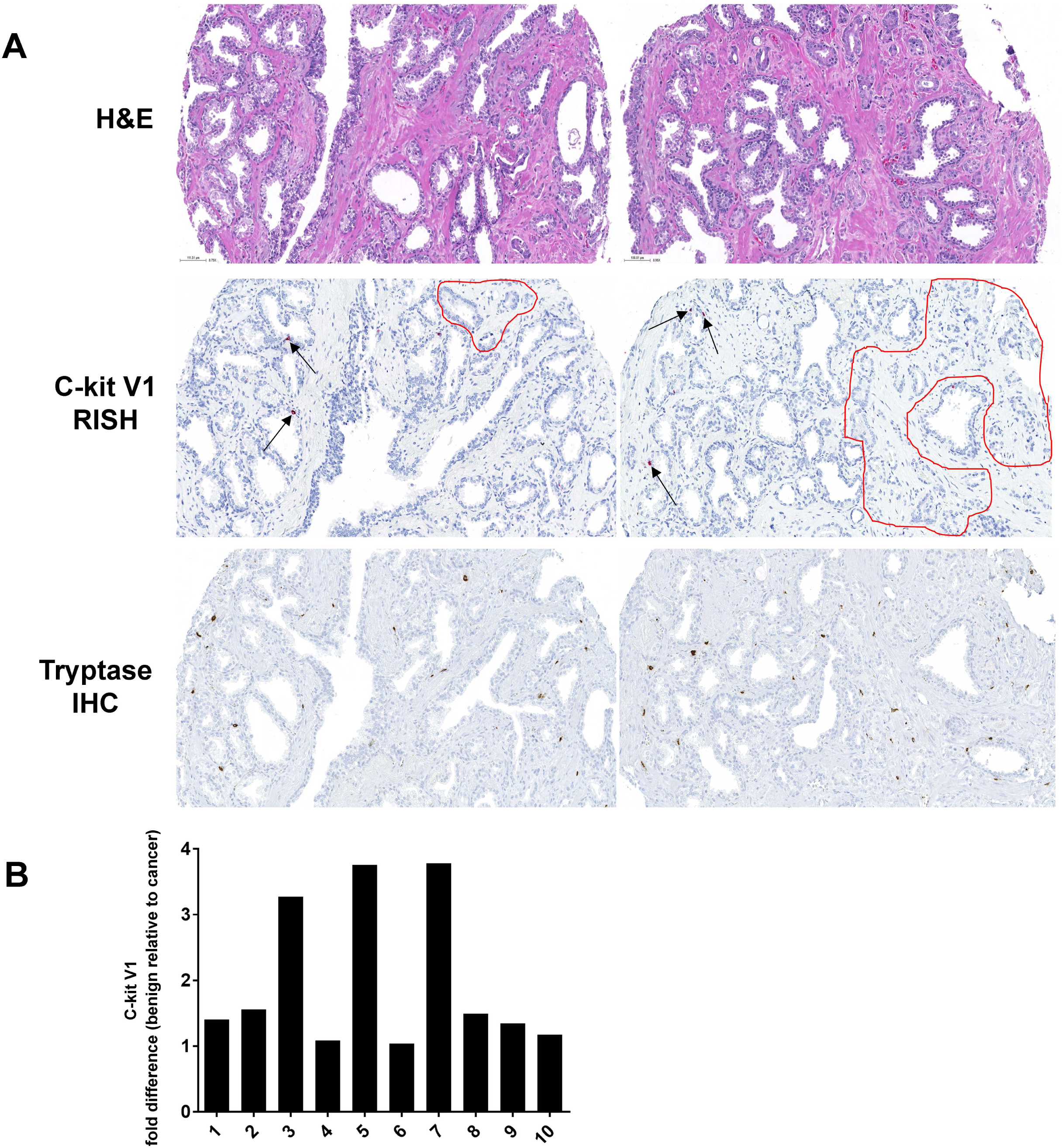
Differential expression of C-kit splice variants in mast cell in benign versus cancer tissues. A) H&E, C-kit V1 RISH, and tryptase IHC (all mast cells) demonstrate association of V1 expressing mast cells in benign versus cancer (red circled area) regions. Only a subset of mast cells were found to express C-kit V1. Sections used for H&E, RISH, and IHC were approximately 10 sections away from each other so the same areas but not the same cells are represented in each assay. B) qRT-PCR for C-kit V1 in RNA isolated from benign versus matched cancer tissues. Shown is the fold difference in C-kit expression normalized to GAPDH in benign relative to cancer.

### Modulation of tumorigenesis by mast cell dosage in the Hi-Myc prostate cancer model

To further explore a functional role for extra-tumoral mast cells in prostate tumorigenesis, we crossed a well-known prostate cancer mouse model, the Hi-Myc mouse, with a mast cell knock-out mouse known as the Wsh mouse. The Hi-Myc mouse is on a BL6 background and over-expresses Myc in an androgen dependent manner in the prostate, developing PIN by 2 weeks and invasive carcinoma in the dorsal lateral prostate by 6 months [37]. Interestingly, we noted that mast cells do not infiltrate into the cancer areas in Hi-Myc mice (Supplemental Figure S6).

Consequently, this model was used to study the contribution of extra-tumoral mast cells, specifically. The Wsh mouse carries an inversion in the Corin promoter in a FVB background mouse, resulting in lack of expression of C-kit and a mast cell knock-out model [38]. Due to the mixed background nature of this cross, Hi-Myc-Wsh mice were taken to the F2 generation to produce littermates of all Myc and Kit genotypes (Hi-Myc +/-, Kit+/+ (WT), Hi-Myc +/-, Kit+/- (HT), Hi-Myc +/-, Kit-/- (KO)) in order to control as much as possible for mixed backgrounds, as well as study the dosage effect of Wsh genotype on mast cell number and tumor invasion.

We began by counting mast cells via toluidine blue stain in each lobe of the prostate from HT and WT mice (KO mice had no mast cells). Interestingly, we found that the dorsolateral lobe (DLP) had the highest number of mast cells of all the lobes in the prostates of both WT and HT mice, but there were fewer mast cells overall in HT mice (Figure 7A). We next quantified the total area of PIN, cribriform PIN, and invasive cancer in each mouse prostate lobe. As expected for the Hi-Myc mouse model, the DLP also preferentially developed invasive cancer compared to the anterior prostate (AP) and ventral prostate (VP, Figure 7B), suggestive that the lobe specificity of invasive caner to the DLP of the Hi-Myc mouse model may be due to the presence of high numbers of mast cells in the dorsolateral lobes of the mice. In further support of this, the total area of invasive cancer was lower in KO and HT mice compared to WT mice, although, interesting only significant for HT mice (p = 0.029, Mann Whitney test, Figure 7C). Likewise, the total area of all involvement with PIN, cribriform PIN (cPIN), and invasive cancer was higher in WT animals than KO and HT mice, although only significant for HT mice (p = 0.023, Figure 7D). These data suggest a dosage effect where higher numbers of extra-tumoral mast cells resulted in higher cancer invasion, further supporting a role for mast cells in tumorigenesis in the Hi-Myc model.

**Figure 7.**
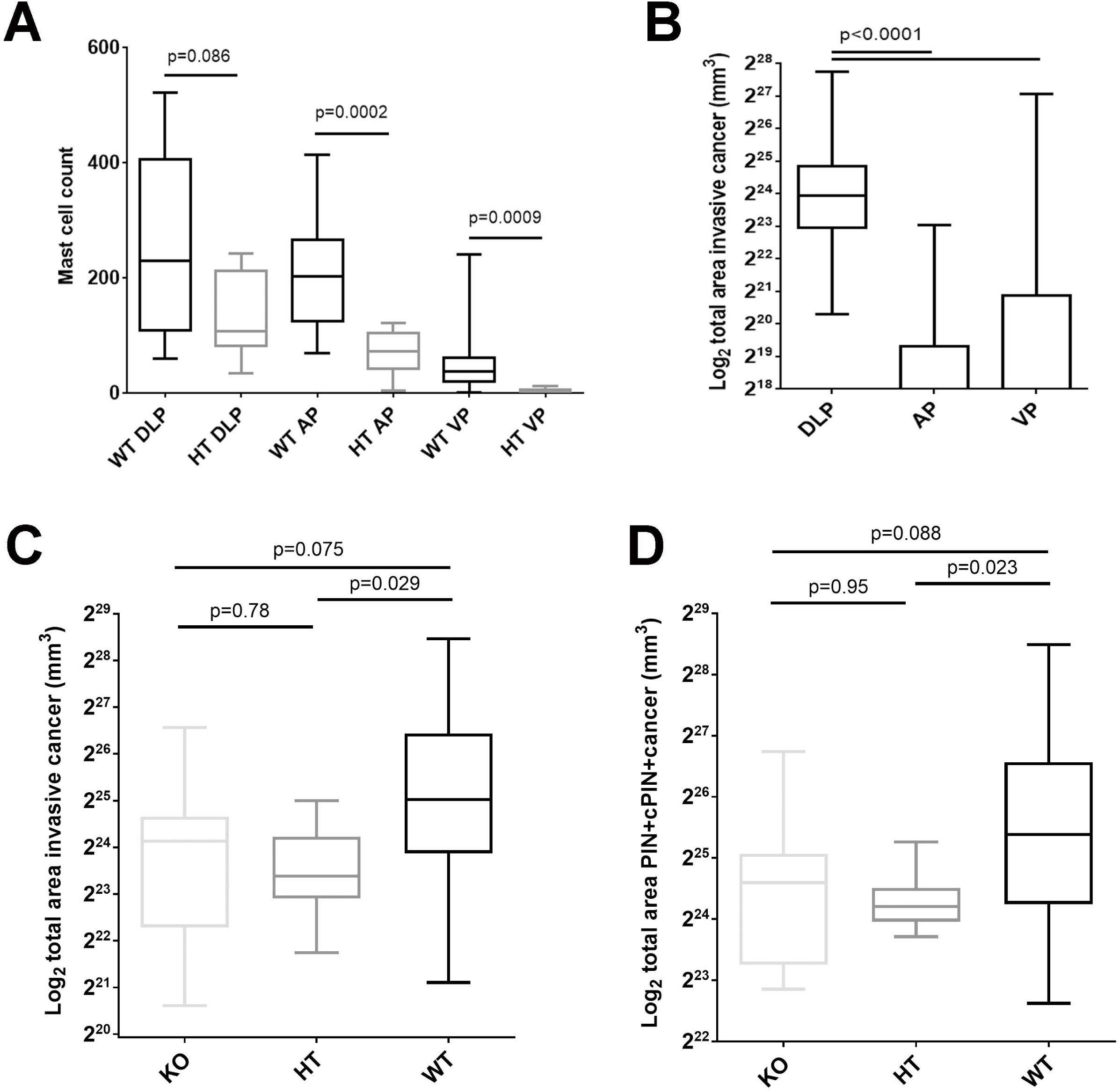
The influence of extra-tumoral mast cells on the Hi-Myc mouse prostate cancer model. A) Mast cell counts in each prostate lobe in mast cell wildtype (WT) and heterozygous (HT) Hi-Myc mice. DLP = dorsolateral, AP = anterior, VP = ventral. B) Log_2_ total invasive area by prostate lobe. C) Log_2_ total invasive cancer area by mouse genotype: mast cell WT, HT, and knockout (KO), n=10 of each genotype. D) Log_2_ total involvement area of PIN, cribriform PIN (cPIN), and invasive cancer by mouse genotype.

## Discussion

Mast cells have been less well studied in the context of the tumor microenvironment than their myeloid cousins such as macrophages. Herein, we have conducted phenotyping of mast cells in prostate cancer extra-tumoral regions as well as within the tumor microenvironment. We began by examining the two classically described mast cell subtypes, MC_T_ and MC_TC_ mast cells, in benign and cancer regions of the prostate an in relation to prostate cancer outcomes. We found that MC_T_ mast cells are more prevalent than MC_TC_ mast cells in both benign and cancer regions, and this trend was consistent among both African-American and European-American men.

Whereas our previous studies have identified an association between high numbers of extra-tumoral mast cells and biochemical recurrence and the development of metastases after radical prostatectomy [14, 24], the present study confirmed this finding and further identified an association with death from prostate cancer as an outcome and that the relationship between extra-tumoral mast cells and prostate cancer outcomes is likely specifically driven by MC_T_ mast cells. Aside from the known differences in their preformed mediators, functional differences in the classical MC_T_ and MC_TC_ mast cell subtypes are not well understood, and this should be an important focus of future studies.

Since the potential significance of mast cell counts varies by whether the mast cells were present in regions of benign tissue or cancer, we further assessed differences in gene expression profiles of benign versus tumor-infiltrating mast cells. We found differential expression of many mast cell and inflammatory pathway-related genes, such as CXCR4 on tumor-infiltrating mast cells. CXCR4 is a chemokine receptor for CXCL12 that is expressed by inflammatory cells including T cells, macrophages, and mast cells, and has been shown to play a role in mast cell recruitment [32, 39, 40]. The CXCL12-CXCR4 pathway in mast cells has been causally implicated in several cancers, including UV induced skin cancer, glioblastoma, and prostate cancer [41-43]. In addition, one study in an obesity and MYC-induced mouse model found that the CXCR4-CXCL12 pathway was a driver in tumor migration and invasion, and that blocking CXCR4 sensitized the mice to chemotherapy [44]. Finally, a previous meta-analysis of human data determined that CXCR4 is significantly more highly expressed in prostate tumor tissue than benign prostate tissue, and is associated with a higher prostate cancer stage, but not Gleason grade [45]. This study also showed an association with lymph node involvement, bone metastasis, and poor prognosis. Our study also found that CXCR4 is more highly expressed in tumor tissues than benign prostate tissue, and using a RISH assay as well as dual-RISH stains we determined that CXCR4 is most highly expressed in cells present within the stromal component of the tumor including mast cells, macrophages, and lymphocytes. Although, our study did find an association between CXCR4 and Gleason grade when we examined higher grade (Grade group 3-5) versus lower grade (Grade group 1-2) prostate cancer.

We likewise noted with interest differential expression of previously characterized variants of C-kit between benign tissue (higher expression of V1) and tumor-infiltrating (higher expression of V2) mast cells. C-kit V1 is expressed in lower levels than V2, especially in the context of cancer [35, 36]. V2 has been shown in multiple studies to be associated with higher colony formation, lower contact inhibition, and higher tumorigenicity in mice when expressed in NIH3T3 fibroblasts [35]. Expression of V2 in early myeloid cells also results in higher and more rapid activation and internalization of SRC family kinases in response to SCF, as well as higher SCF-dependent growth and a stronger chemotactic response to SCF in-vitro compared to early myeloid cells expressing V1 [46]. In addition, V2 has been associated with higher levels of granule formation, histamine content, and growth in mast cells as well as in faster response to SCF compared to V1, however V1 expression resulted in longer activation [47, 48]. Finally, a higher serum V2/V1 ratio has also been associated with higher levels of neoplastic mast cells in mastocytosis [47]. Thus, mast cells expressing V2 are demonstrably more active than those expressing V1, which we find intriguing in light of the fact that loss of intra-tumoral mast cells is associated with worse outcomes. We postulate that mast cells within the tumor may participate in anti-tumor immunity whereas extra-tumoral mast cells may serve a different role that paradoxically drives tumorigenesis.

Finally, we examined the influence of extra-tumoral mast cell numbers on tumor invasion and growth in the Hi-Myc mouse model of prostate cancer. We found that mast cells do not infiltrate Hi-Myc tumors, which is analogous to our previous finding that, with the exception of bone metastases, human prostate cancer metastases are devoid of intra-tumoral mast cells [24]. Using this model, we studied the effect of extra-tumoral mast cells numbers on carcinogenesis. We noted with interest that HT mice (with decreased numbers of mast cells compared to WT mice) developed significantly less invasive cancer and pre-invasive lesions. The mechanism of this complex relationship will certainly be the focus of future studies.

In conclusion, our study indicates that there may be multiple different mast cell subtypes present within the prostate, and that mast cells present in extra-tumoral regions are not only distinct from those infiltrating prostate cancer, but they may also hold an important prognostic significance in terms of adverse prostate cancer outcomes.

## Acknowledgements

This work was supported by Department of Defense Prostate Cancer Research Program, Award Numbers W81XWH-14-1-0364 (to H.H.S., A.M.D. and K.S.S.) and W81XWH-17-1-0286 (to T.L.L., C.M.H., C.E.J. and K.S.S.) and W81XWH-18-2-0013 (K.S.S.) and W81XWH-18-2-0015 (A.M.D.) Prostate Cancer Biorepository Network (PCBN), Prostate Cancer Foundation Challenge Award 19CHAS03 (J.P.M., A.M.D., T.L.L., and K.S.S.). We would like to thank Dr. Srinivasan Yegnasubramanian, Dr. Sarah Wheelan, and the members of the SKCCC Next Generation Sequencing Core and the OTS Core, supported by NCI grant P30CA006973, for assistance with RNAsequencing and tissue microarray construction, respectively. We also thank Dr. Charles Drake for helpful advice and discussion.

## Supplemental Figure Legends

**Supplemental Figure S1.**
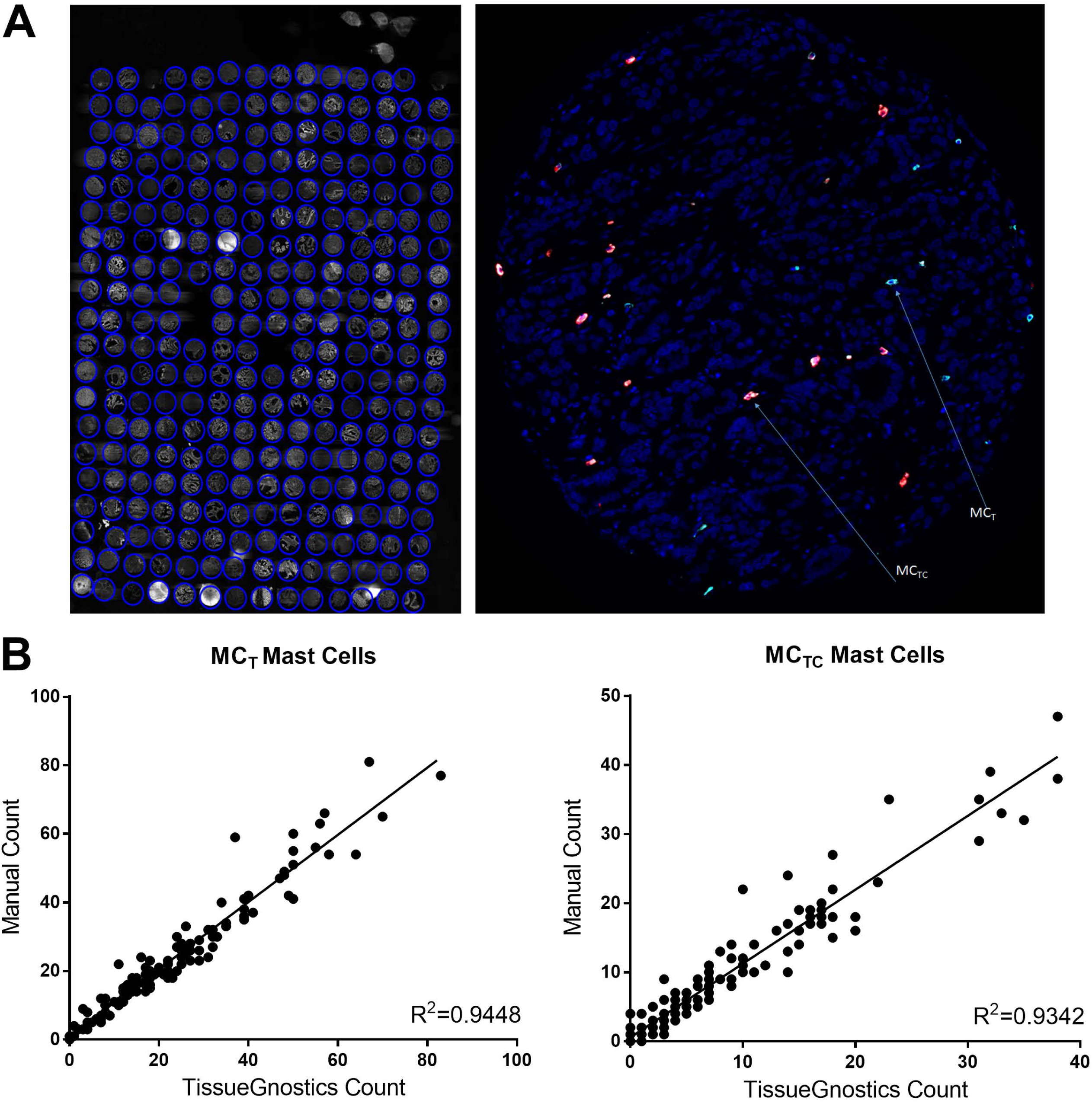
A) Dual IF for mast cell tryptase (green) and chymase (red). Left, TMA visualized by IF. Right, individual TMA spot showing mast cell subtypes visualized using dual IF, where MC_TC_ were tryptase and chymase double positive cells (arrow denoted as MC_TC_), and MC_T_ were tryptase only positive cells (arrow denoted as MC_T_). B) Manual counting of mast cell subtypes versus automated counting using TissueGnostics software.

**Supplemental Figure S2.**
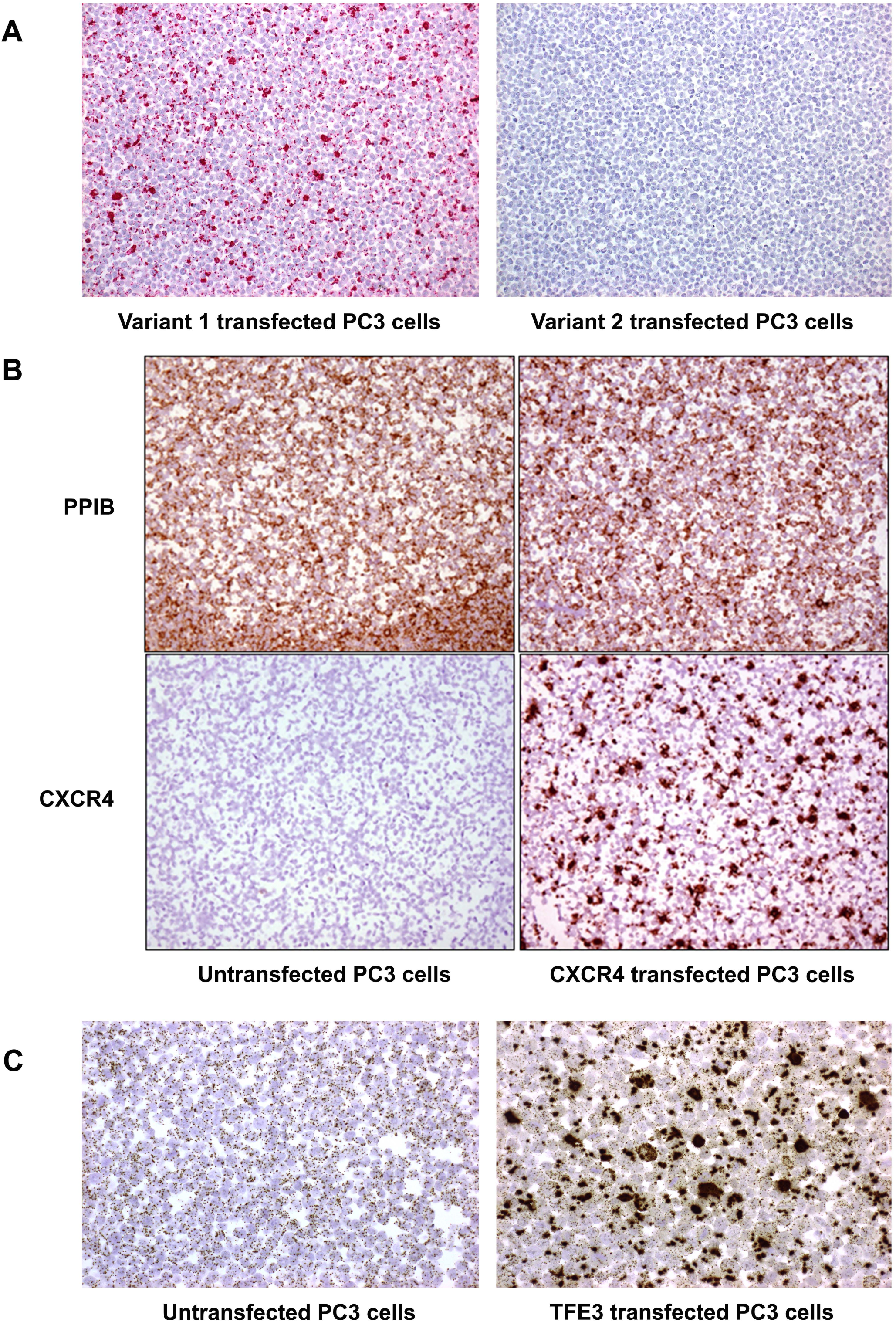
Positive and negative controls for C-kit V1, C-kit V2, TFE3, and CXCR4 RISH. A) PC3 cells transfected with c-Kit variant 1 or C-kit variant 2 expression vectors and stained with a Basescope c-Kit variant 1-specific RISH probe (red). B) Untransfected or CXCR4-transfeted PC3 cells stained with PPIB (positive control) or CXCR4 RISH (brown). C) Untransfected or TFE3-transfeted PC3 cells stained with TFE3 RISH (brown). PC3 cells were found to express basal levels of TFE3 that was markedly increased upon transfection with the TFE3 expression vector.

**Supplemental Figure S3.**
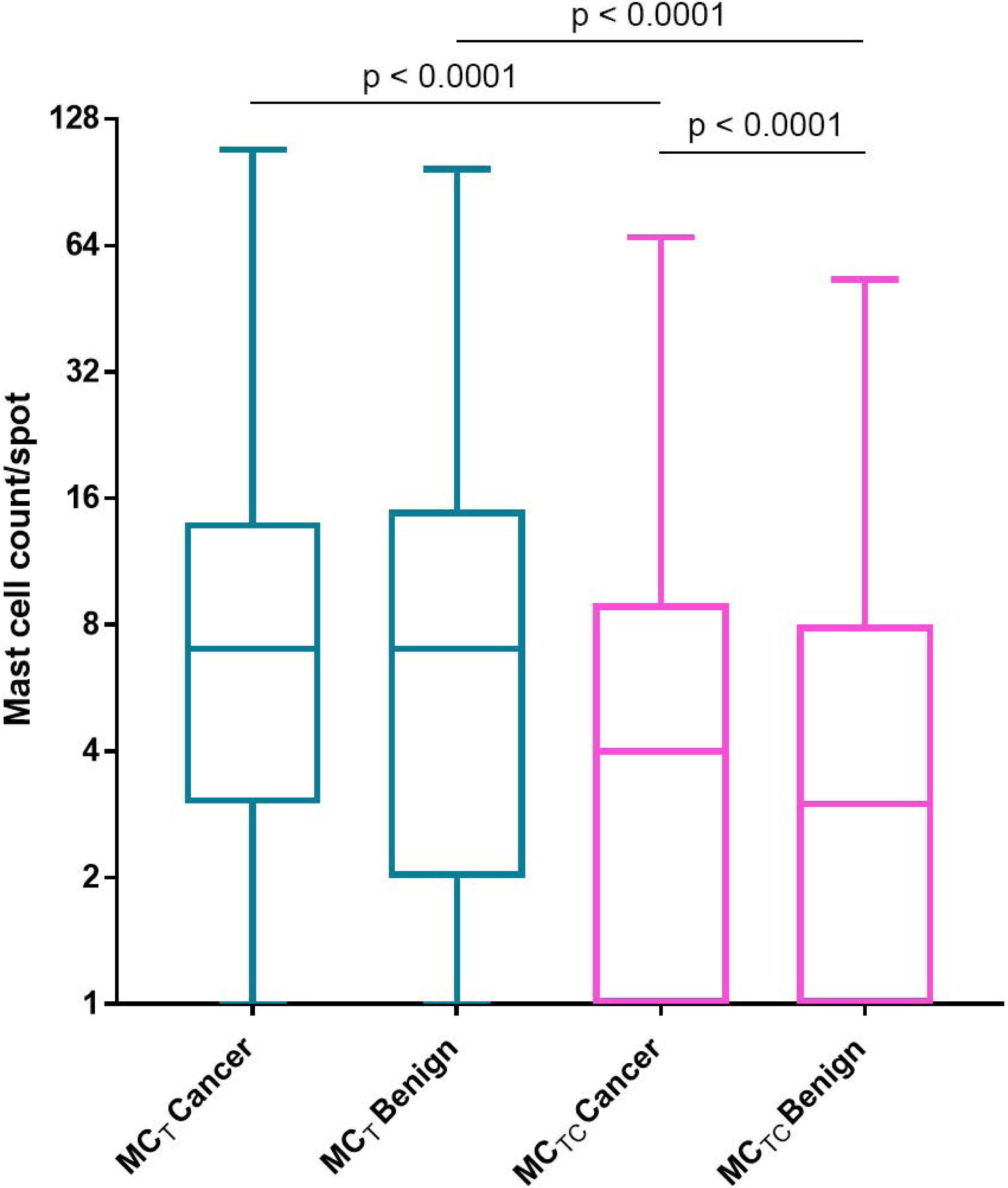
Log2 of mast cell count per tissue spot of each mast cell subtype in benign and cancer tissue spots in the Intermediate/High Risk TMA set.

**Supplemental Figure S4.**
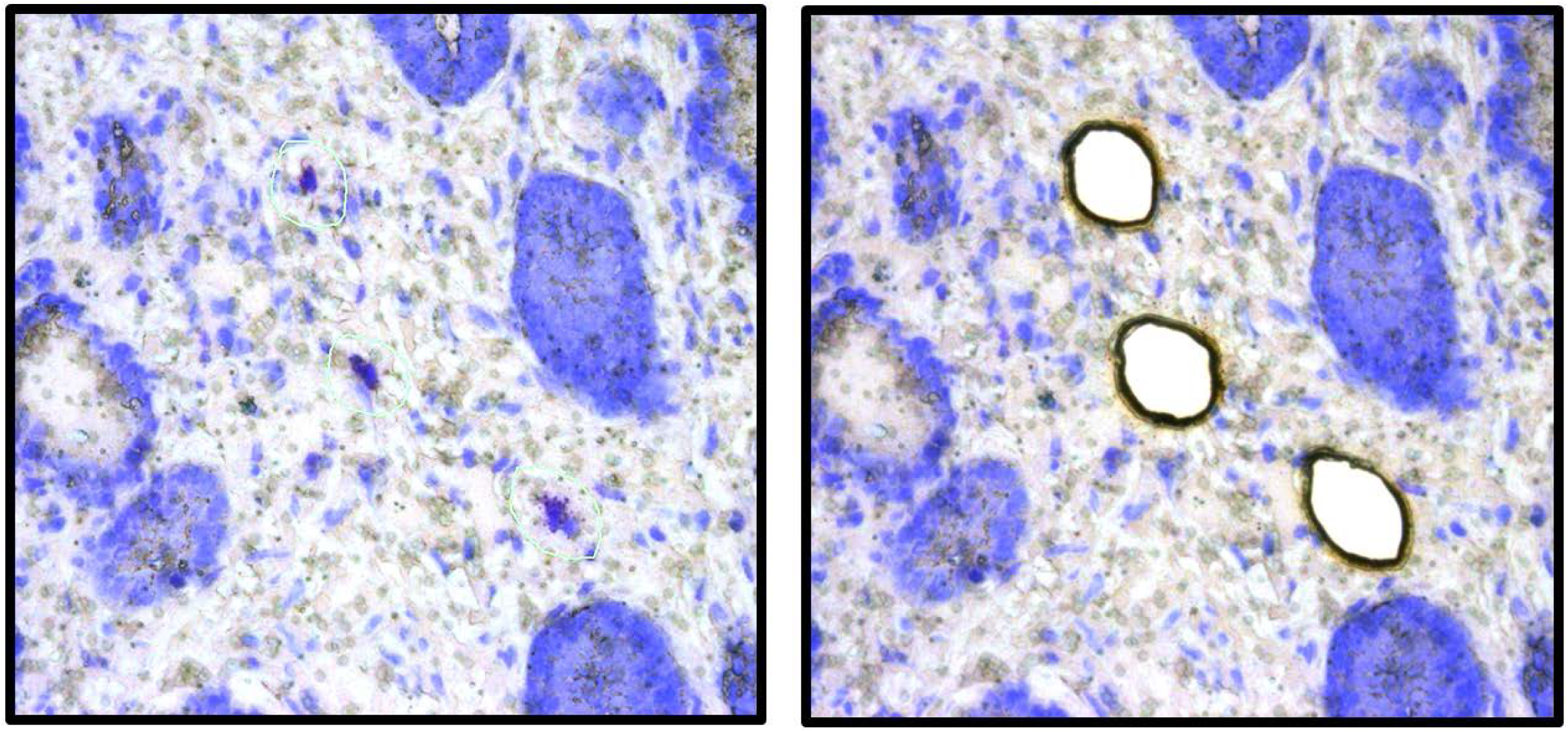
Example of LCM of toluidine blue stained mast cells from prostate tissue.

**Supplemental Figure S5.**
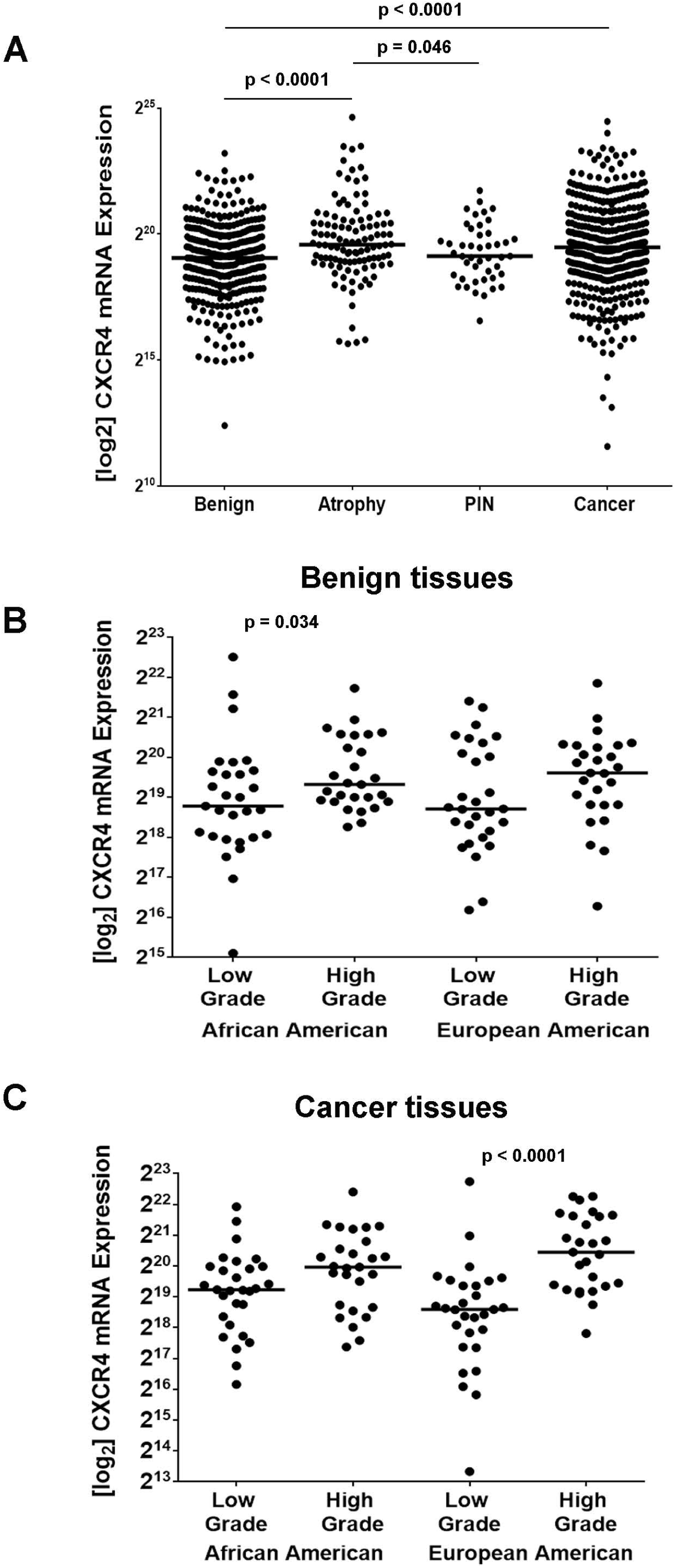
Quantification of CXCR4 expression in the 120 Case High Grade Race TMA set. A) Log_2_ CXCR4 expression per TMA spot in TMA spots that were benign, contained atrophy, contained PIN, or were cancer. B) Log_2_ of median CXCR4 expression per case in benign tissues in low grade (Grade group 1-2) versus higher grade (Grade group 3-5) cases in African American versus European American men. C) Log_2_ of median CXCR4 expression per case in cancer tissues in low grade (Grade group 1-2) versus higher grade (Grade group 3-5) cases in African American versus European American men.

**Supplemental Figure S6.**
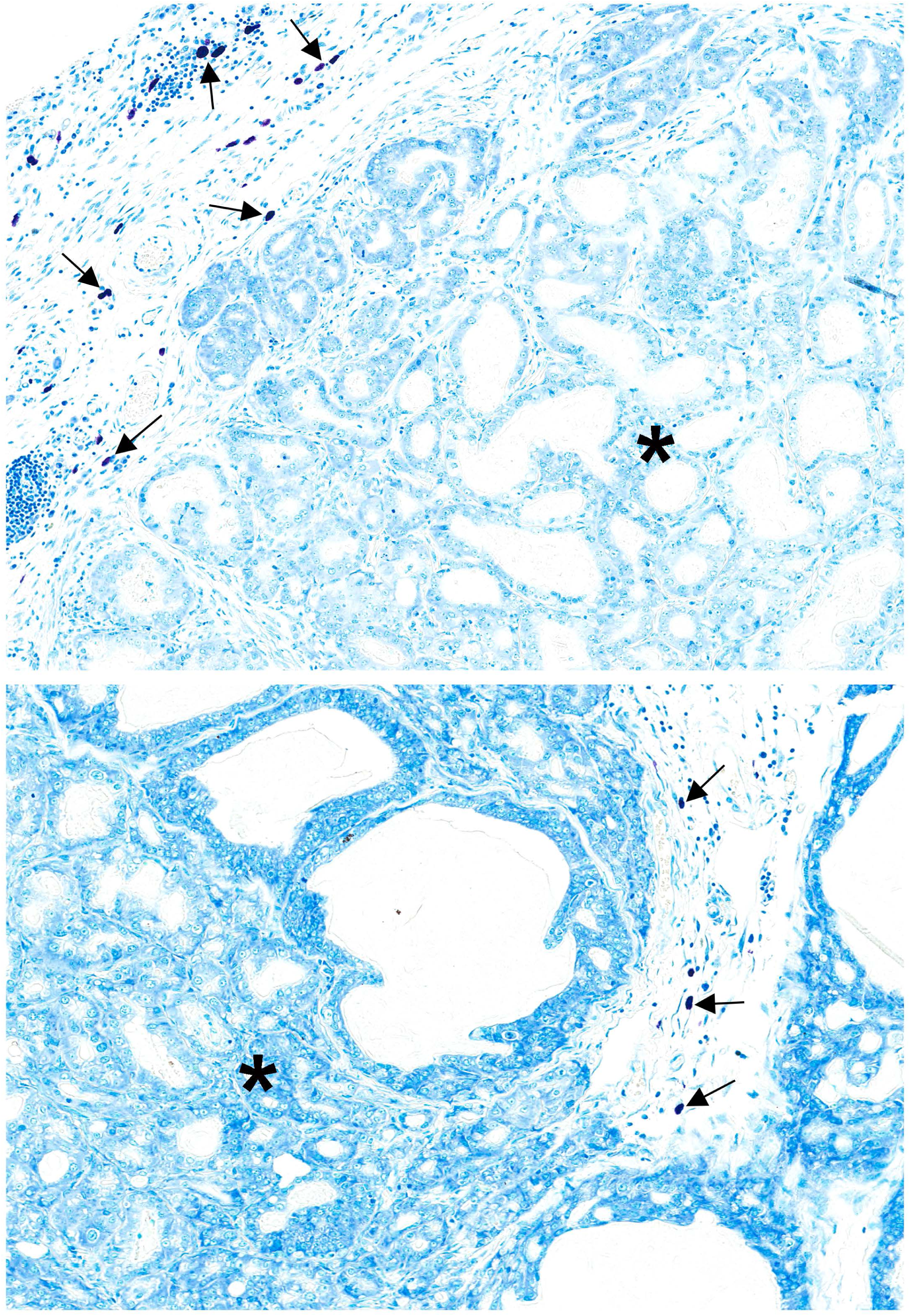
Toluidine blue stain of WT Hi-Myc mouse prostate. Mast cells are present in extra-tumoral stromal regions (arrows), but are not present within the tumors (asterisks).

## Supplemental Methods

### RNAseq library prep and analysis

Whole transcriptome RNA amplification was performed for pooled RNA from tumor and benign tissue mast cells with the Nugen Ovation RNA-seq System v2 kit using barcodes (Tecan, Männedorf, Switzerland). RNAseq libraries were sequenced on an Illumina HS2500 Rapid Run 100bp x 100bp paired end. Illumina’s CASAVA 1.8.2 was used to convert BCL files to FASTQ files. Default parameters were used. rsem-1.2.29 was used for running the alignments as well as generating gene and transcript expression levels [1]. The ‘rsem-calculate-expression’ module was used with the following options:

--bowtie-chunkmbs 200

--calc-ci

--output-genome-bam

--paired-end

--forward-prob

The data was aligned to “hg19” reference genome. rsem-1.2.29’s EBSeq was used for Differential Expression analysis [2]. ‘rsem-for-ebseqfind-DE’ was used to run EBSeq, default parameters were used.

## References

1. Amin K. The role of mast cells in allergic inflammation. Respiratory Medicine 2012;106:9–14.

2. Fajt ML, Wenzel SE. Mast cells, their subtypes, and relation to asthma phenotypes. Annals of the American Thoracic Society 2013;10:S158–S64.

3. da Silva EZM, Jamur MC, Oliver C. Mast cell function: a new vision of an old cell. J Histochem Cytochem 2014;62:698–738.

4. Balzar S, Fajt ML, Comhair SA, et al. Mast cell phenotype, location, and activation in severe asthma. Data from the Severe Asthma Research Program. American journal of respiratory and critical care medicine 2011;183:299–309.

5. Johansson A, Rudolfsson S, Hammarsten P, et al. Mast cells are novel independent prognostic markers in prostate cancer and represent a target for therapy. Am J Pathol 2010;177:1031–41.

6. Andersson CK, Bergqvist A, Mori M, et al. Mast cell-associated alveolar inflammation in patients with atopic uncontrolled asthma. The Journal of allergy and clinical immunology 2011;127:905–12.e1-7.

7. Méndez-Enríquez E, Hallgren J. Mast cells and their progenitors in allergic asthma. Frontiers in Immunology 2019;10.

8. Galinsky DST, Nechushtan H. Mast cells and cancer—No longer just basic science. Critical Reviews in Oncology/Hematology 2008;68:115–30.

9. Lee Y-M, Jippo T, Kim D-K, et al. Alteration of protease expression phenotype of mouse peritoneal mast cells by changing the microenvironment as demonstrated by in situ hybridization histochemistry. The American Journal of Pathology 1998;153:931–6.

10. Gurish MF, Pear WS, Stevens RL, et al. Tissue-regulated differentiation and maturation of a v-abl-immortalized mast cell-committed progenitor. Immunity 1995;3:175–86.

11. Sfanos K, Hempel H, De Marzo A. The role of inflammation in prostate cancer, In: Aggarwal BB, Sung B, Gupta SC, (eds). Inflammation and Cancer. Vol. 816. Springer Basel: 2014. pp 153–81.

12. Varricchi G, Galdiero MR, Loffredo S, et al. Are mast cells MASTers in cancer? Frontiers in immunology 2017;8:424-.

13. Rajput AB, Turbin DA, Cheang MCU, et al. Stromal mast cells in invasive breast cancer are a marker of favourable prognosis: a study of 4,444 cases. Breast Cancer Research and Treatment 2008;107:249–57.

14. Hempel HA, Cuka NS, Kulac I, et al. Low intratumoral mast cells are associated with a higher risk of prostate cancer recurrence. The Prostate 2017;77:412–24.

15. Fleischmann A, Schlomm T, Kollermann J, et al. Immunological microenvironment in prostate cancer: high mast cell densities are associated with favorable tumor characteristics and good prognosis. Prostate 2009;69:976–81.

16. Siiskonen H, Poukka M, Bykachev A, et al. Low numbers of tryptase+ and chymase+ mast cells associated with reduced survival and advanced tumor stage in melanoma. Melanoma research 2015;25:479–85.

17. Glajcar A, Szpor J, Pacek A, et al. The relationship between breast cancer molecular subtypes and mast cell populations in tumor microenvironment. Virchows Archiv 2017;470:505–15.

18. Carlini MJ, Dalurzo MC, Lastiri JM, et al. Mast cell phenotypes and microvessels in non-small cell lung cancer and its prognostic significance. Hum Pathol 2010;41:697–705.

19. Fleischmann A, Schlomm T, Köllermann J, et al. Immunological microenvironment in prostate cancer: High mast cell densities are associated with favorable tumor characteristics and good prognosis. The Prostate 2009;69:976–81.

20. Globa T, Saptefrti L, Ceausu RA, et al. Mast cell phenotype in benign and malignant tumors of the prostate. Polish Journal of Pathology 2014;65:147–53.

21. Taverna G, Giusti G, Seveso M, et al. Mast Cells as a Potential Prognostic Marker in Prostate Cancer. Disease markers 2013;35:711–20.

22. Sari, Serel, Çandir Ö, Öztürk, Kosar. Mast cell variations in tumour tissue and with histopathological grading in specimens of prostatic adenocarcinoma. BJU International 1999;84:851–3.

23. Johansson A, Rudolfsson S, Hammarsten P, et al. Mast cells are novel independent prognostic markers in prostate cancer and represent a target for therapy. The American Journal of Pathology 177:1031–41.

24. Hempel Sullivan H, Heaphy CM, Kulac I, et al. High extratumoral mast cell counts are associated with a higher risk of adverse prostate cancer outcomes. Cancer epidemiology, biomarkers & prevention : a publication of the American Association for Cancer Research, cosponsored by the American Society of Preventive Oncology 2020;29:668–75.

25. Globa T, Saptefrti L, Ceausu RA, et al. Mast cell phenotype in benign and malignant tumors of the prostate. Polish journal of pathology : official journal of the Polish Society of Pathologists 2014;65:147–53.

26. Aslan A, Erdem H, Balta H, et al. Tryptase and Chymase Expression Differences in Prostatic Adenocarcinomas. Sakarya Medical Journal 2018;8:229–34.

27. Ellwood-Yen K, Graeber TG, Wongvipat J, et al. Myc-driven murine prostate cancer shares molecular features with human prostate tumors. Cancer Cell 2003;4:223–38.

28. Baena Del Valle J, Zheng Q, Hicks J, et al. Rapid loss of RNA detection by in situ hybridization in stored tissue blocks and preservation by cold storage of unstained slides. American Journal of Clinical Pathology 2017;In Press.

29. Ross AE, Johnson MH, Yousefi K, et al. Tissue-based genomics augments post-prostatectomy risk stratification in a natural history cohort of intermediate- and high-risk men. European urology 2016;69:157–65.

30. Darshan M, Zheng Q, Fedor HL, et al. Biobanking of derivatives from radical retropubic and robot-assisted laparoscopic prostatectomy tissues as part of the prostate cancer biorepository network. Prostate 2014;74:61–9.

31. Iwata T, Schultz D, Hicks J, et al. MYC overexpression induces prostatic intraepithelial neoplasia and loss of Nkx3.1 in mouse luminal epithelial cells. PLoS One 2010;5:e9427.

32. Juremalm M, Hjertson M, Olsson N, et al. The chemokine receptor CXCR4 is expressed within the mast cell lineage and its ligand stromal cell-derived factor-1alpha acts as a mast cell chemotaxin. European journal of immunology 2000;30:3614–22.

33. Juremalm M, Nilsson G. Chemokine receptor expression by mast cells. Chemical immunology and allergy 2005;87:130–44.

34. Yagil Z, Hadad Erlich T, Ofir-Birin Y, et al. Transcription factor E3, a major regulator of mast cell–mediated allergic response. Journal of Allergy and Clinical Immunology 2012;129:1357–66.e5.

35. Caruana G, Cambareri AC, Ashman LK. Isoforms of c-KIT differ in activation of signalling pathways and transformation of NIH3T3 fibroblasts. Oncogene 1999;18:5573–81.

36. Pedersen M, Ronnstrand L, Sun J. The c-Kit/D816V mutation eliminates the differences in signal transduction and biological responses between two isoforms of c-Kit. Cellular signalling 2009;21:413–8.

37. Ellwood-Yen K, Graeber TG, Wongvipat J, et al. Myc-driven murine prostate cancer shares molecular features with human prostate tumors. Cancer cell 2003;4:223–38.

38. Nigrovic PA, Gray DH, Jones T, et al. Genetic inversion in mast cell-deficient (Wsh) mice interrupts corin and manifests as hematopoietic and cardiac aberrancy. The American journal of pathology 2008;173:1693–701.

39. Minami H, Nagaharu K, Nakamori Y, et al. CXCL12-CXCR4 Axis Is Required for Contact-Mediated Human B Lymphoid and Plasmacytoid Dendritic Cell Differentiation but Not T Lymphoid Generation. Journal of immunology (Baltimore, Md : 1950) 2017;199:2343–55.

40. Niu Q, Zhou Q, Liu Y, Jiang H. Expression of CXCR4 on T-cell subsets and Plasma IL-17 Concentrations in Patients with Aplastic Anaemia. Sci Rep 2017;7:9075.

41. Ellem SJ, Taylor RA, Furic L, et al. A pro-tumourigenic loop at the human prostate tumour interface orchestrated by oestrogen, CXCL12 and mast cell recruitment. The Journal of pathology 2014;234:86–98.

42. Polajeva J, Sjosten AM, Lager N, et al. Mast cell accumulation in glioblastoma with a potential role for stem cell factor and chemokine CXCL12. PloS one 2011;6:e25222.

43. Sarchio SNE, Scolyer RA, Beaugie C, et al. Pharmacologically antagonizing the CXCR4-CXCL12 chemokine pathway with AMD3100 inhibits sunlight-induced skin cancer. The Journal of investigative dermatology 2014;134:1091–100.

44. Saha A, Ahn S, Blando J, et al. Proinflammatory CXCL12-CXCR4/CXCR7 Signaling Axis Drives Myc-Induced Prostate Cancer in Obese Mice. Cancer research 2017;77:5158–68.

45. Chen Q, Zhong T. The association of CXCR4 expression with clinicopathological significance and potential drug target in prostate cancer: a meta-analysis and literature review. Drug design, development and therapy 2015;9:5115–22.

46. Young SM, Cambareri AC, Odell A, Geary SM, Ashman LK. Early myeloid cells expressing c-KIT isoforms differ in signal transduction, survival and chemotactic responses to Stem Cell Factor. Cellular signalling 2007;19:2572–81.

47. Chan EC, Bai Y, Bandara G, et al. KIT GNNK splice variants: expression in systemic mastocytosis and influence on the activating potential of the D816V mutation in mast cells. Experimental hematology 2013;41:870–81.e2.

48. Montero JC, Lopez-Perez R, San Miguel JF, Pandiella A. Expression of c-Kit isoforms in multiple myeloma: differences in signaling and drug sensitivity. Haematologica 2008;93:851–9.

## References

1. Li B, Dewey CN. RSEM: accurate transcript quantification from RNA-Seq data with or without a reference genome. BMC Bioinformatics 2011;12:323.

2. Leng N, Kendziorski C. EBSeq: An R package for gene and isoform differential expression analysis of RNA-seq data. R package version 1.24.0. 2019.

